# A closed translocation channel in the substrate-free AAA+ ClpXP protease diminishes rogue degradation

**DOI:** 10.1101/2022.08.27.505532

**Authors:** Alireza Ghanbarpour, Steven E. Cohen, Xue Fei, Tristan A. Bell, Tania A. Baker, Joseph H. Davis, Robert T. Sauer

## Abstract

Intracellular proteases must be specific to avoid degrading the wrong proteins. Here, we present cryo-EM structures of *E. coli* ClpXP, a AAA+ protease, which reveal that the axial channel of ClpX is closed prior to the binding and subsequent translocation of a protein substrate. An open-channel ClpX mutation stimulates degradation of casein, a non-specific substrate, indicating that channel closure contributes to increased degradation specificity. We demonstrate that ClpX activates ClpP cleavage of a degron-free decapeptide by a channel-independent mechanism, in which the peptide substrate appears to pass through a symmetry mismatched gap in the interface between ClpX and ClpP before entering the degradation chamber via the axial portal of ClpP. The peptide products of ClpXP protein degradation are likely to exit the chamber by the reverse route.

## INTRODUCTION

AAA+ proteases carry out ATP-dependent protein degradation in all cells^1^. In a series of energy-consuming reactions, a AAA+ ring hexamer binds a target protein, unfolds any native structure present, and translocates the unfolded polypeptide though an axial channel and into the proteolytic chamber of a peptidase for degradation. ClpXP, a well-characterized AAA+ protease, consists of one or two copies of the ClpX_6_ unfoldase/translocase and the double-ring ClpP_14_ peptidase^2,3^. Access to the ClpP chamber is controlled by axial portals in each heptamer, which are restricted in free ClpP to limit degradation of unfolded polypeptides, but open to a diameter of ∼30 Å when ClpX binds^4-9^. Robust degradation is facilitated by non-specific contacts between ClpX and translocating polypeptides, ensuring that any sequence can be spooled into ClpP, and by a channel through the ClpX ring that connects to an axial portal of ClpP and thus to the proteolytic chamber^7-10^. However, unlike Lon and many other AAA+ proteases, ClpXP does not degrade proteins unfolded by heat shock or other cellular stresses^11-12^. Given the common structural features and mechanisms of AAA+ proteases^13^, this higher level of ClpXP substrate specificity is intriguing.

In *Escherichia coli* and other proteobacteria, ClpXP plays a critical role in protein-quality control by degrading substrates with C-terminal ssrA tags or degrons, which are generated by tmRNA-mediated ribosome rescue when protein biosynthesis stalls^14-15^. The *E. coli* ssrA tag ends in an Ala-Ala sequence, which is critical for ClpX recognition of ssrA-tagged proteins and additional cellular substrates^16-17^. In a cryo-EM structure called a recognition complex (pdb 6WRF), these C-terminal residues of the ssrA tag bind in the upper portion of the ClpX axial channel, the remainder of which is blocked by a pore loop of one ClpX subunit^8^. In another structure (pdb 6WSG), the ssrA tag moves down into an open ClpX channel to enable the ATP-dependent mechanical reactions necessary for substrate translocation and degradation^8^.

Is the axial channel of substrate-free ClpX closed or does ssrA-tag binding stabilize channel closure? Here, by determining cryo-EM structures of substrate-free ClpXP, we show that the channel is closed in the same fashion as in the substrate-bound recognition complex. Importantly, we also demonstrate that mutation of the pore loop that closes the ClpX channel increases the rate of degradation of a non-specific substrate. Finally, we present experiments that illustrate how ClpX stimulates ClpP cleavage of non-specific peptides as well as the inverse mechanism by which the peptide products of protein degradation are released from the chamber of ClpXP.

## RESULTS

### Cryo-EM structures of substrate-free ClpXP have a closed translocation channel

We determined three cryo-EM structures of *E. coli* ClpXP without protein substrate at resolutions ranging from 2.6 to 3.1 Å (**Figures 1, S1-S5**; **Table S1**). These structures were very similar (0.3-0.9 Å pairwise Cα RMSDs) despite differences that included: (*i*) whether complexes contained full-length ClpX or ^SC^ClpX^ΔN^ (a single-chain pseudohexamer with genetically tethered subunits lacking the N-domain); (*ii*) whether ClpXP complexes were stabilized with ATP or ATPγS; and (*iii*) whether structures were solved using doubly capped (ClpX_6_•ClpP_14_•ClpX_6_) or singly capped (ClpX_6_•ClpP_14_) particles (**Table S1**). ^SC^ClpX^ΔN^ was used for prior cryo-EM studies of *E. coli* ClpXP and supports ClpP degradation of ssrA-tagged substrates^8-9,18^. For the structures that used doubly capped ClpXP particles, only one of the two ClpX rings was included in the final refinement mask as one hexamer was always poorly resolved relative to the other. This likely occurs because the conformations of the two ClpX hexamers are uncoordinated, and thus the resulting density map of one ClpX hexamer represents a errant average of multiple conformations.

**Figure 1.**
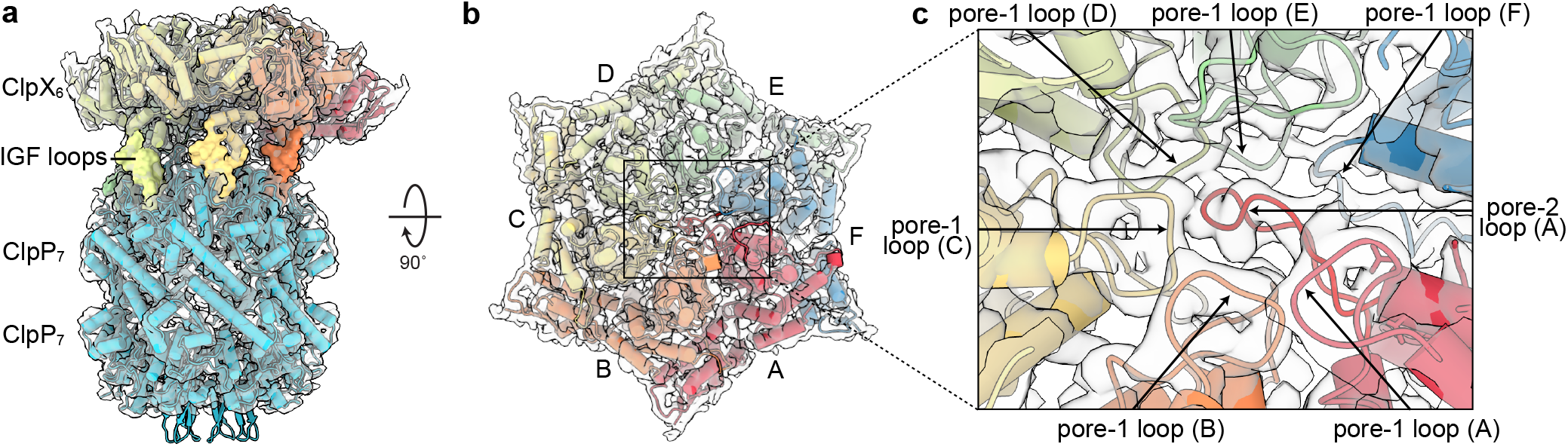
Cryo-EM structure of substrate-free ^SC^ClpX^ΔN^•ClpP particles. Side (**a**) and top (**b**) views of ^SC^ClpX^ΔN 6^•ClpP_14_ structure. Cryo-EM density is colored light grey and fitted atomic models are displayed in cartoon representation and colored and labeled by ClpX subunit. Key structural features are noted. Inset (**c**) of the ClpX axial pore highlights the locations of pore-1 and pore-2 loops from each subunit.

The N-terminal domains of ClpX were not visible in the full-length structure (**Figure S4, S5**), presumably as a consequence of their flexible linkage to the AAA+ body of ClpX. The final models typically included residues 65-413 of each ClpX subunit, four ATPγS/ATP and two ADP molecules, and residues 2-193 of each ClpP subunit. Residues 192-203 of the pore-2 loop of the lowest subunit in the ClpX spiral (chain F) had poor density and were not modeled. The axial portal of the distal Clp ring in the singly capped structure was closed (**Figure S5**), whereas the axial portals of ClpP rings contacting ClpX were open.

Cryo-EM studies of full-length *Listeria monocytogenes* ClpXP, which had been chemically crosslinked, reported interactions between two ClpX hexamers, mediated by their N-terminal domains^6^. We saw no evidence of similar interactions in micrographs, 2D class averages, or the final density map for our full-length *E. coli* ClpXP dataset.

The substrate-free structures aligned well with the substrate-bound recognition complex (less than 1 Å Cα RMSDs). Moreover, in the structures without protein substrate, the pore-2 loops of ClpX subunit A blocked the translocation channel as observed previously in the recognition complex (**Figures 1, 2**). Thus, channel closure is not induced by ssrA binding. By contrast, the axial channels in other *E. coli* or *Neisseria meningitidis* ClpXP structures with translocating substrates are open, and the pore-2 loop of ClpX subunit A is partially disordered^7-9^.

**Figure 2.**
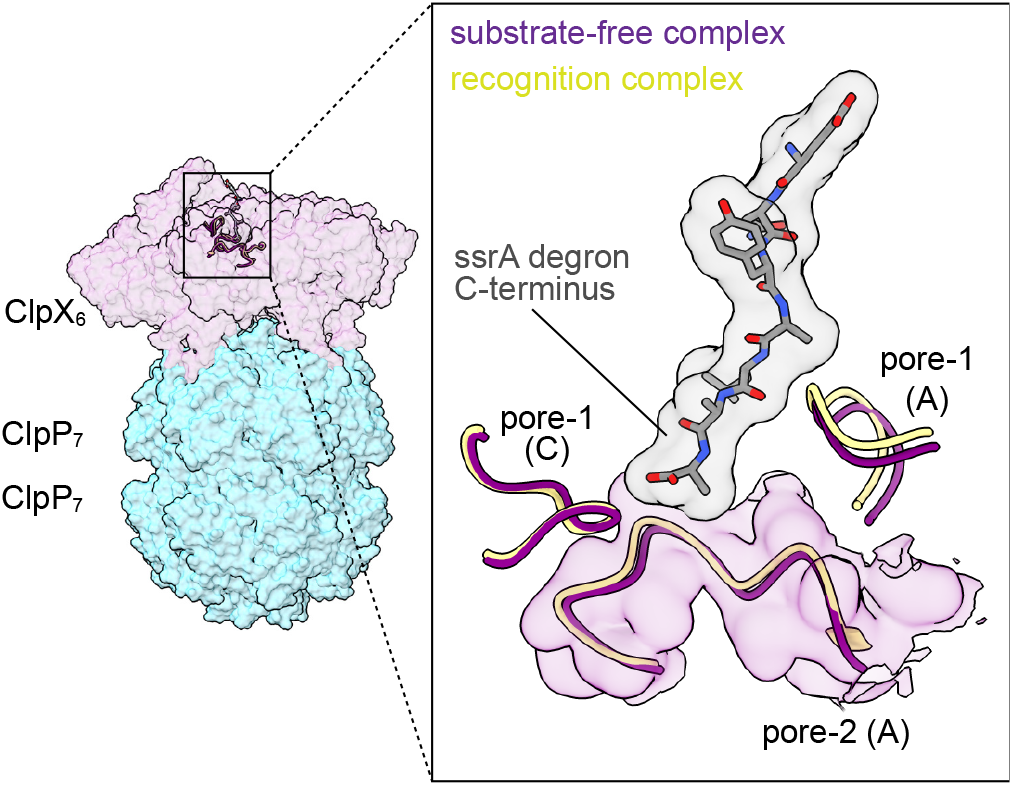
ClpX pore-1 and pore-2 loops adopt similar conformations in substrate-free and recognition complexes. Side view of the ^SC^ClpX^ΔN^•ClpP complex with semi-transparent surface representation of ClpX (purple) and ClpP (cyan). Inset highlights similar conformations of pore-1 (residues 150-156) and pore-2 (residues 195-205) loops from the subunits A and C in the substrate-free (purple) and recognition complex (yellow) structures. An atomic model of the ssrA degron from the recognition complex is depicted as a stick model and gray surface representation.

### Multistep ssrA-tag binding

The RKH loop of ClpX subunit C in our substrate-free structures adopted a conformation that would clash with the ssrA degron (**Figure 3**), suggesting a model in which the tag binds in two steps. In this model, the C-terminal Leu-Ala-Ala-_coo_^−^ of the ssrA tag initially binds the pore-1 loops of ClpX chains A and B and the pore 2 loop of chain A in essentially the same conformation as observed in our substrate-free structures, possibly displacing the RKH loop of chain C. In a second step, illustrated in the morph in **Supplementary Movie 1**, downward movement of the RKH loop of subunit C then stabilizes the bound conformation of the ssrA degron, as observed in the recognition complex.

**Figure 3.**
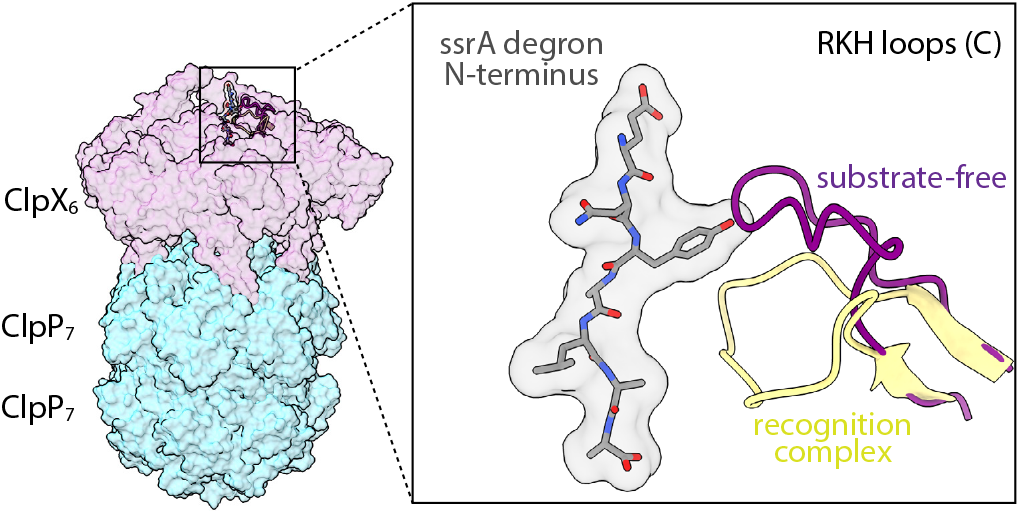
Flexible RKH loops are in altered conformations in the presence and absence of substrate. The structure on the left shows a semi-transparent surface representation of the ClpXP substrate free ^SC^ClpX^ΔN^•ClpP complex. The inset highlights the ssrA degron (stick representation with transparent surface overlayed) from the ClpXP recognition complex and RKH loops of chain C from both structures (residues 220-238; cartoon representation). Note that the aromatic Tyr side chain in the ssrA tag would sterically impinge on the substrate-free conformation (purple) of this RKH loop.

A power stroke, initiated from the recognition complex, can open the ClpX channel and move the substrate polypeptide six residues deeper into the pore^8^. ClpX hydrolyzes ATP in the absence of protein substrates^19^, and thus there may be stages during the ATPase cycle when the channel opens transiently. Using cryoDRGN^20^, we inspected volumes generated at 100 *k*-means cluster centers, and identified only one volume with an open channel. Conservatively, we thus estimated that less than 10% of the ClpX in our substrate-free dataset had an open channel conformation.

### The closed translocation channel enhances degradation specificity

‘Clp’ stands for caseinolytic protease, but ClpXP degrades the non-specific casein protein very slowly^21-23^. To test if an open-channel variant degraded casein more rapidly, we used ^pore2Δ/ins^ClpX^ΔN^ in which pore-2 loop residues 191-201 were replaced with a Gly_2_-Ser_2_-Gly_2_ sequence^24^, as we hypothesized that this variant cannot make the interactions that stabilize the blocked-channel conformation. Notably, the degradation rate of FITC-casein by ^pore2Δ/ins^ClpX^ΔN^•ClpP was ∼5-fold faster than the rate of degradation by ClpX^ΔN^•ClpP (**Figure 4**). Hence, mutational opening of the channel enhances degradation of a non-specific substrate.

**Figure 4.**
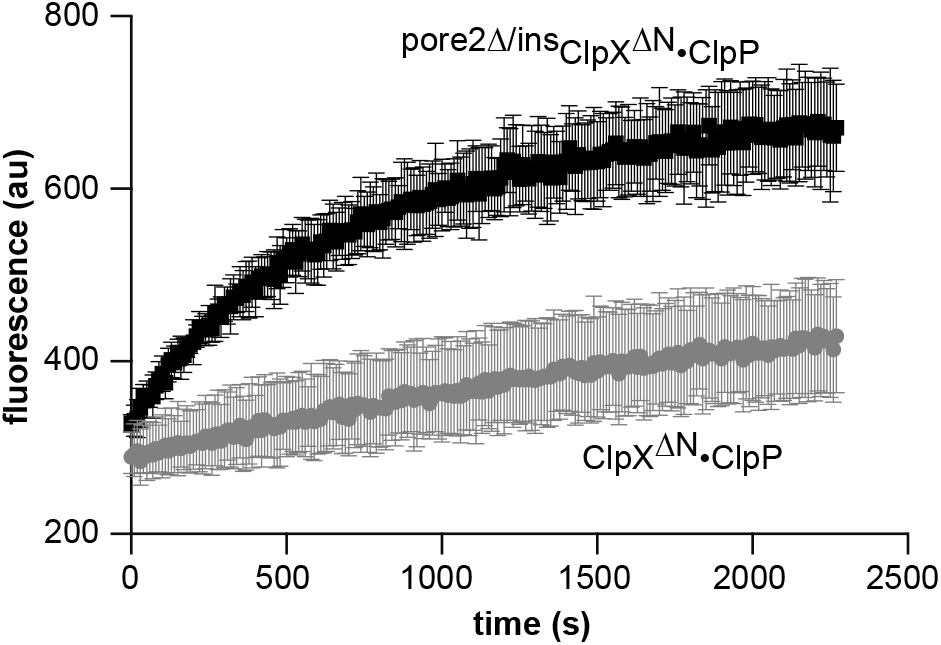
Degradation of casein by ClpX•ClpP. Mutation of the ClpX pore-2 loop enhances the rate of FITC casein degradation, as determined by decreased auto-quenching of FITC fluorescence upon ClpP proteolysis. Reaction contained FITC casein (50 µM), ^pore2Δ/ins^ClpX^ΔN^ or ^SC^ClpX^ΔN^ (1 µM) and ClpP (20 µM) (see Methods for detailed conditions). Symbols denote mean fluorescence (n = 3), and error bars represent ± 1 SD.

### ClpXP cleavage of small peptides implies a pathway for product release

The axial portals of ClpP control entry of substrates into the degradation-chamber. These portals are closed in ClpP alone and in the distal Clp ring of our singly capped structure (**Figure S5**) but open when in direct contact with ClpX^4,6-9,25^. This conformational change accounts for the ∼10-fold stimulation of ClpP cleavage of a nonspecific decapeptide by ClpX^26^. However, the closed channel in our substrate-free structures is inconsistent with decapeptides entering ClpP in the same fashion as protein substrates. We tested the hypothesis that a decapeptide need not pass through the ClpX channel to access the ClpP portal by assaying decapeptide cleavage in the presence or absence of *E. coli* DHFR-_gsylaalaa_, which is a good ClpXP substrate^27^ and should thus fill the channel. Importantly, the rate of decapeptide cleavage decreased only marginally when near saturating concentrations of DHFR-_gsylaalaa_ were present (**Figure 5**), both when ATP-dependent protein degradation by ClpX^ΔN^**•**ClpP was ongoing and when degradation by ^E185Q^ClpX^ΔN^**•**ClpP was prevented by the E^185^Q Walker-B mutation that dramatically slows ATP hydrolysis but does not impair substrate binding^28^. Thus, decapeptide cleavage does not require transit through the ClpX channel.

**Figure 5.**
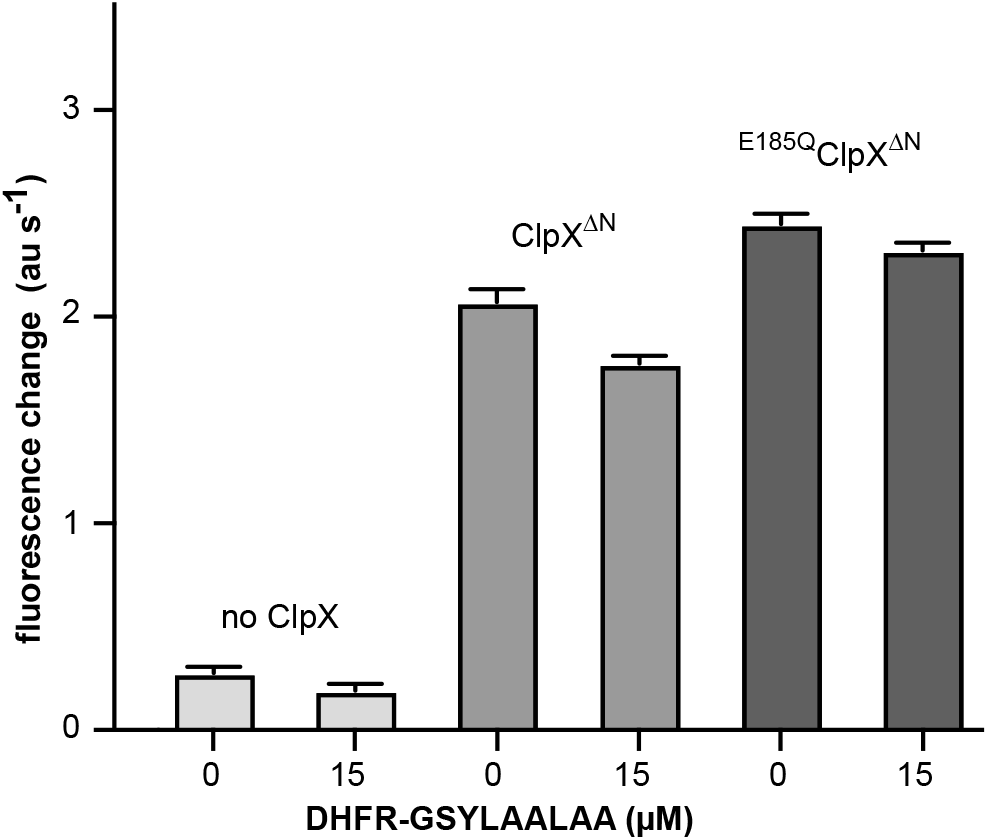
ClpP-mediated decapeptide cleavage. Rates of ClpP cleavage of Abz-KASPVSLGY^NO2^D (15 µM) were assayed by increased fluorescence in the presence/absence of ClpX variants (1 µM), ClpP (50 nM), and the indicated concentrations of a DHFR-_gsylaalaa_ protein substrate. Values are means (n = 3-6) ± 1 SD. Notably, in the presence of ClpX variants, the DHFR substrate only marginally reduced the rate of decapeptide cleavage.

In the presence of the 20-fold excess of ClpX^ΔN^ used in this experiment, most ClpP tetradecamers were doubly capped and the axial portal of the distal ClpP ring in singly capped enzymes was closed (**Figure S5**). How then do decapeptides enter the degradation chamber of ClpXP? We propose that they first pass through the asymmetric interface between ClpX and ClpP. Six flexible IGF loops from an *E. coli* ClpX hexamer dock into six of seven hydrophobic clefts in a ClpP heptamer, leaving one unoccupied cleft (**Figure 6**). Thus, peptides could pass between the IGF loops of ClpX that span the empty cleft and then through the open ClpP portal, obviating the need to traverse the ClpX channel. This ‘IGF-gap’ pathway might be even more accessible if only five of the IGF loops were stably docked, as observed in one structure of *N. meningitidis* ClpX^ΔN^**•**ClpP^9^. Moreover, interactions between the IGF loops of *E. coli* ClpX and ClpP are dynamic^29^, creating possibilities for additional or widened gaps.

**Figure 6.**
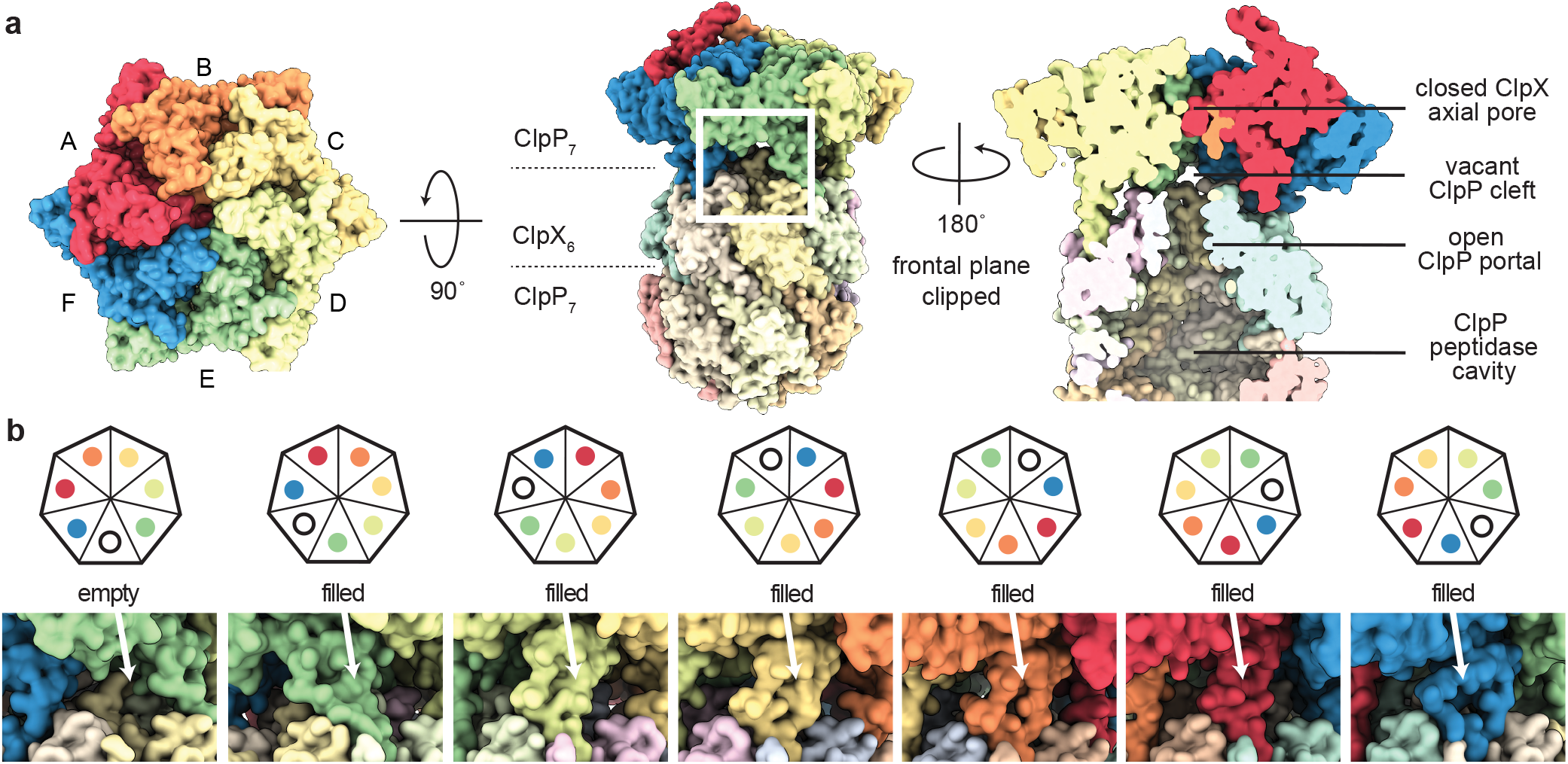
A symmetry mismatch at the ClpX•ClpP interface may allow peptide product release. (**a**) Top-view (left) and side-view (middle) surface representations of the ^SC^ClpX^ΔN^•ClpP complex. Surface representation clipped in-plane at the ClpXP midline (right) with key structural features noted. (**b**) Schematic representation of ClpX (colored circles) and ClpP (heptagon) subunit interfaces (top) and surface representation of the modeled ^SC^ClpX^ΔN^•ClpP complex highlighting each ClpP cleft (bottom). Subunits are colored as in panel (a) and labeled to indicate the occupancy status of the highlighted ClpP cleft.

A related question is how the peptide products of ClpXP cleavage, which typically range from 5 to 20 residues long^30-31^, exit the degradation chamber following cleavage of a substrate polypeptide? It has been proposed that structural fluctuations at the equatorial ring-ring interface of ClpP create transient gaps or windows that allow product egress^32^. Our results suggest another mechanism. Notably, if peptides can enter the degradation chamber of ClpXP by passing between gapped IGF loops while protein degradation is ongoing, then the principle of microscopic reversibility dictates that peptide products of similar size produced by ClpP degradation, could exit the ClpP chamber by the same pathway. Importantly, the ∼30-Å diameter of the open ClpP portal is wide enough to accommodate an incoming translocating polypeptide from ClpX, while simultaneously allowing outgoing peptide products to exit via the same portal.

### How do other AAA+ proteases regulate specificity?

If a closed translocation channel of ClpX is biologically important in preventing rogue protein degradation, then how do other AAA+ proteases deal with the same problem? At low temperature, the axial channel of the *E. coli* HslUV protease is also blocked, albeit by a completely different molecular mechanism. Specifically, regions of a domain unique to the AAA+ HslU hexamer form a trimeric plug that engages and blocks access to the translocation channel^33^. This autoinhibitory plug melts at higher temperatures, activating proteolysis by HslUV. In substrate-free structures of the human 26S proteasome, the translocation channel of the AAA+ unfolding ring is open but the axial portal into the degradation chamber is typically closed^34^, providing another mechanism to limit non-specific degradation. Substrate-free structures of Lon protease from *E. coli* and *Yersinia pestis* form open lock-washer spirals^35-36^. How these structures bind substrate and transition to the active closed-spiral conformation is presently unclear, although the rates of refolding of denatured substrates and the interplay between rescue by chaperones and proteolysis by Lon may determine the specificity of this protease^37^. Substrate-free structures of the ClpAP protease have not been reported, but in biochemical experiments ClpAP binds acid-denatured GFP and degrades casein rapidly, whereas wild-type ClpXP has neither of these activities^22,38^, suggesting that the axial channel of ClpAP is open in the absence of substrates. ClpAP specificity in the cell might be enhanced by adapter proteins but this model remains to be tested.

## MATERIALS AND METHODS

### Protein purification

C-terminally His^6^-tagged variants of *E. coli* ClpP, *E. coli* ClpX^ΔN^ (residues 62-424), and *E. coli* ^SC^ClpX^ΔN^ (six ClpX^ΔN^ subunits connected by genetic tethers) were expressed separately in *E. coli* and purified as described^39-40^. *E. coli* ClpP with a C-terminal His_6_ tag and full-length ClpX (with a neutral K408E mutation) encoded in pT7 plasmids were expressed in *E. coli* strain HMS174(DE3) and purified as reported^41^. After protein expression at 30 °C for 3-4 h, cells were harvested by centrifugation at 4,500 rpm for 30 min, resuspended in 50 mM Tris HCl (pH 8.0), 100 mM KCl, 1 mM MgCl_2_, 5 mM DTT, and 10% glycerol, and lysed using a cell disruptor. Insoluble material was removed by centrifugation at 25,000 rpm in a Ti45 rotor for 30 min. ClpX in the supernatant fraction was purified using ammonium-sulfate precipitation (35%), phenyl-Sepharose chromatography (Amersham HP 17-1082-01), and chromatography on a self-packed mono-Q ion-exchange column. His_6_-tagged ClpP was purified using Ni^++^-NTA affinity and size-exclusion chromatography^39^. The ^pore2Δ/ins^ClpX^ΔN^ variant was expressed and purified as described^24^. Purified proteins were concentrated, flash frozen in liquid nitrogen, and stored at -80 °C.

### Cryo-EM sample preparation

For the substrate-free ^SC^ClpX^ΔN^•ClpP structure, ^SC^ClpX^ΔN^ (3 µM pseudohexamer) and ClpP (1.5 µM tetradecamer) in 20 mM HEPES (pH 7.5), 100 mM KCl, and 25 mM MgCl_2_ were incubated with ATP (5 mM) for 15 min at room temperature to ensure that protein substrate present as a consequence of copurification was degraded. Prior to vitrification, samples (2 µL) were placed on 400-mesh Quantifoil™ 2/1 copper grids, which had been glow-discharged for 60 s in an easiGlow glow discharger (Pelco) at -15 mA, and were blotted using a FEI Vitrobot Mk IV instrument for 4 s with a blot force of +10 (22 °C; 99% relative humidity). The full-length substrate-free structure was obtained from an experiment in which wild-type ClpX and ClpP were incubated at room temperature with ATPΥS (2.5 mM) for 5 min at room temperature in EM buffer [25 mM HEPES (pH 7.5), 100 mM KCl, and 25 mM MgCl_2_], prior to addition of λΟ-Αrc and ATPΥS. Final concentrations were ClpX_6_ (5.8 µM), ClpP_14_ (1.9 µM), λΟ-Αrc (20 µM dimer; this substrate protein was absent from the structure solved), and ATPΥS (2.5 mM). Samples (2.5 µL) were placed on 200-mesh Quantifoil™ R 2/1 copper grids, which had been glow-discharged for 60 s in an easiGlow glow discharger (Pelco) at -15 mA, and were blotted using a FEI Vitrobot Mk IV instrument for 3 s with a blot force of +0 (6 °C; 99% relative humidity) and immediately vitrified in liquid ethane.

### Cryo-EM data collection

For the substrate-free ^SC^ClpX^ΔN^•ClpP structure, 2,405 movies were collected with EPU (Thermo Fisher Scientific) on a Titan Krios G3i (Thermo Fisher Scientific), operating at an acceleration voltage of 300 kV and magnification of 105,000 X and detected in super resolution mode on a K3 detector (Gatan) with an effective pixel size of 0.87 Å (0.435 Å super resolution). Movies were collected as 30 frames with a defocus range from -0.75 to -2.5 µm and a total exposure per specimen of 49.4 e-/Å^2^. For the full length ClpXP structure, 11,719 movies were collected with EPU (Thermo Fisher Scientific) using aberration-free image shift (AFIS) and hole-clustering method on a Titan Krios G3i (Thermo Fisher Scientific) with an acceleration voltage of 300 kV and magnification of 130,000 X and detected in super-resolution mode on a K3 detector (Gatan) for an effective pixel size of 0.679 Å (binned by 2). Movies were collected as 40 frames with a defocus range from -0.3 to -1.75 µm and a total exposure per specimen of 52.7 e^−^/Å^2^.

### Cryo-EM pre-processing and particle picking

For the substrate-free ^SC^ClpX^ΔN^•ClpP structure, data processing was performed using cryoSPARC (v.3.3.1) and default parameters unless noted^44^. Raw movies (2,405) were pre-processed using ‘*Patch motion correction*’, and ‘*Patch CTF estimation*’ jobs. Visual inspection revealed that particles primarily adopted ‘top’ and ‘side’ views. Approximately 150 such side views were manually selected with picks centered on the ClpX•ClpP interface and particles were extracted (box size 440 × 440 pixels, Fourier cropped to 320 × 320 pixels) and averaged into a single class using the ‘*2D classification*’ utility. Using this 2D class average, we applied the ‘*Template picker*’ to 20 micrographs, performed a round of 2D classification on the resulting 2,254 particles, and selected two representative side-view classes from the resulting 20 classes. These classes were then used for template picking of the full dataset. ‘*Exposure curation*’ was used to eliminate 252 micrographs with poor CTF fits and/or too few particles. Extraction of side views from the remaining 2,153 micrographs resulted in 154,554 particles (box size 440 × 440 pixels) that were Fourier-cropped to 320 × 320 pixels. A similar workflow was applied to isolate top views, relying on a ‘*Blob picker’* (minimum particle diameter 40; max particle diameter 150; use ring blob; min separation distance 4) instead of the manual picker used in the initial stage, resulting in 237,063 top-view particles. Side view and top view particles were subjected to ‘*2D classification*’ and curation by selecting high resolution classes, resulting in 140,011 and 227,251 particles, respectively. These were then combined, and the resulting set was subjected to the ‘*Remove duplicate particles*’ utility, resulting in a preliminary stack of 333,774 particles. The explicitly singly capped ^SC^ClpX^ΔN^•ClpP structure used a subset of this particle stack as described below.

Data processing for the full-length ClpXP structure was performed using cryoSPARC (v.3.3.1) and default parameters unless noted^44^. Raw movies (11,709) were pre-processed using ‘*Patch motion correction*’, and ‘*Patch CTF estimation*’. 171,780 particles were picked using the blob-picker tool through 1,000 random micrographs. Particles were extracted (box size 440 × 440 pixels, Fourier cropped to 256 × 256 pixels) and averaged using the ‘*2D classification*’ utility. Using this 2D class average, we then applied the ‘*Template picker*’to the full dataset and extracted the resulting particles (box size 484 × 484 pixels, Fourier cropped to 320 × 320 pixels). After another round of ‘*2D classification’*, 1,903,710 particles were selected. The micrographs were subjected to ‘*Manually curate exposures*’, which excluded 30 micrographs based on CTF estimated resolution and particle count. Particles were then extracted (box size 440 × 440 pixels, Fourier cropped to 256 × 256 pixels), and subjected to ‘*2D classification*’. 1,801,963 particles were selected from these 2D classes as a preliminary stack.

### *Ab-initio* reconstruction, global refinement, and model building

*Ab-initio* reconstruction and homogeneous refinement using substrate-free ^SC^ClpX^ΔN^•ClpP particles generated a map of the complex at ∼4 Å resolution. To better center particles on a single ClpX•ClpP interface, these particles were re-extracted using translations from this reconstruction and a larger box (initial box 600 × 600 pixels; Fourier cropped to 256 × 256 pixels) and subjected to 2-class ‘*Ab-initio reconstruction’*, which produced an isolated ClpP volume (146,991 particles) and a clear ^SC^ClpX^ΔN^•ClpP volume (175,043 particles). Particles corresponding to the ^SC^ClpX^ΔN^•ClpP volume were then refined to a GSFSC resolution of 3.2 Å, and particle alignments were used for a final round of particle re-extraction (initial box 680 × 680 pixels; Fourier cropped to 256 × 256 pixels), ‘*Ab-initio reconstruction’* and ‘*Homogeneous refinement*’ with per-particle defocus optimization, resulting in a GSFSC ∼2.9 Å map. Using this map, a mask corresponding to the ClpX and the cis ring of ClpP was generated in Chimera X (1.3)^45^ and, following a terminal round of ‘*Duplicate particle removal*’ the resulting 171,869 curated, centered particles were subjected to ‘*Local refinement*’ in cryoSPARC using the aforementioned mask, resulting in a 3.1 Å GSFCS map focused on ^SC^ClpX^ΔN^ and the proximal ClpP ring.

For the ^SC^ClpX^ΔN^•ClpP structure determined specifically from singly capped particles, the initial models and particles resulting from the two class *ab-initio* reconstruction described above were used in to two class ‘*Heterogeneous refinement*’. Particles within the class most closely resembling singly capped ^SC^ClpX^ΔN^•ClpP were then subjected to ‘*Homogeneous refinement*’, producing a ∼4 Å map of the complex. The particles were re-extracted from this reconstruction with a larger box centered on clpP (initial box 900 pixels; down-sampled to 300 pixels) and subjected to ab initio, homogeneous refinement, and duplicate particle removal, resulting in 138,629 particles. These particles were used in 4-class ‘*Ab-initio reconstruction’*. Only one class that corresponded to singly capped ^SC^ClpX^ΔN^•ClpP was selected. Further ‘*Homogeneous refinement*’ of these 49,824 particles yielded a map at 3.1 Å GSFSC resolution.

For the full-length ClpX•ClpP structure, ‘*Ab-initio reconstruction’* was performed using four classes. One class containing the majority of particles (952,541) was selected for ‘*Homogeneous refinement*’ with per-particle defocus optimization; the remaining classes represented junk particles, free ClpP, or a low resolution ClpXP complex. After homogeneous refinement, particles were recentered around the ClpX-ClpP interface. After another round of ‘*Homogeneous refinement*’, ‘*Heterogeneous refinement’* was performed using four classes. A class that contained the majority of particles (527,257) was selected for further ‘*Homogeneous refinement*’ with per-particle defocus optimization. These particles were recentered on the ClpX-ClpP interface and were classified using a three class ‘*Heterogeneous refinement*’ that used a low-pass filtered map resulting from a ‘*Homogeneous refinement’* of the centered 527,257 particle stack. The best resolved class (217,969 particles) was selected and subjected to further ‘*Homogeneous refinement*’ and ‘*Local refinement*’ using a mask focused on ClpX and the cis ClpP ring. The final full-length ClpXP map had a GSFSC resolution of 2.6 Å.

To correct for pixel size errors, the final maps were rescaled using calibrated pixel sizes (0.654 Å and 0.416 Å for 130,000 X and 105,000 X magnifications), which were derived by analyzing the real space correlation between maps of apo-ferritin collected at MIT.nano under identical imaging conditions to those used here with publicly available apo-ferritin density maps.

After map validation using global and local resolution^46^ estimation tools in cryoSPARC, and model building was performed using a combination of ChimeraX (1.3)^45^, Coot (0.9.4)^46^, and Phenix (1.14)^47^. Final maps were sharpened using Phenix for model building, but are unsharpened in all figures.

### Biochemical Assays

Casein degradation was performed at 30 °C using 50 µM FITC-casein (C0528; Sigma Aldrich), 1 µM of ^pore2Δ/ins^ClpX^ΔN^ or otherwise isogenic ClpX^ΔN^, and 20 µM of ClpP in 25 mM HEPES (pH 7.5), 100 mM KCl, 10 mM MgCl_2_, 1 mM DTT, 4 mM ATP, 5 mM phosphocreatine, and 0.05 mg/ml creatine kinase. The concentration of FITC casein was measured using a NanoDrop spectrometer (ThermoFischer Scientific) at an absorbance of 280 nm (ε = 11,460 M^−1^ cm^−1^), and the concentrations of ClpX and ClpP were calculated using extinction coefficients obtained from https://web.expasy.org/protparam. All components except casein were initially mixed at 30 °C in a 384-well assay plate (Corning, 3575) for 15 min. FITC-casein, which had also been preincubated at 30 °C, was then added to initiate degradation, which was monitored by increases in fluorescence (excitation 360 nm; emission 525 nm with a cutoff of 515 nm) using a SpectraMax M5 plate reader.

Cleavage of a synthetic Abz-KASPVSLGY^NO2^D decapeptide (where Abz is a 2-aminobenzoic acid fluorophore and Y^NO2^ is a 3-nitrotyrosine quencher)^25^ by ClpP (50 nM) was assayed in the presence or absence of ClpX^ΔN^ or ^E185Q^ClpX^ΔN^ (1 µM) and with or without 15 µM *E. coli* ^D132C/C152S^DHFR-_gsylaalaa_ with an N-terminal H^6^ and AviTag sequence^42^. The fluorescence of this decapeptide (excitation 320 nm; emission 420 nm) increases following ClpP cleavage. Assays were performed at 37 °C in buffer containing 25 mM HEPES (pH 7.6), 100 mM KCl, 20 mM MgCl_2_, 1 mM EDTA, 10% glycerol, 5 mM ATP, 32 mM creatine phosphate (Roche), and 0.08 mg/mL creatine kinase (Millipore-Sigma).

## ACKNOWLEDGMENTS

Supported by NIH grants R01-GM144542, R35-GM141517 and 5T32-GM007287, and NSF-CAREER grant 2046778.

Samples were prepared at the Automated Cryogenic Electron Microscopy Facility in MIT.nano and screened on a Talos Arctica microscope, which was a gift from the Arnold and Mabel Beckman Foundation.

## CONFLICTS OF INTEREST

The authors declare no conflicts of interest.

## CONTRIBUTIONS

AG, TA Bell, and XF purified proteins and performed biochemical assays. AG, SEC and XF prepared samples for EM imaging and collected data. AG and JHD processed EM data and performed reconstruction and refinement. AG and RTS built and refined the models. All authors contributed to writing and editing the manuscript. TA Baker, JHD, and RTS supervised the project.

## SUPPLEMENTARY FIGURES

**Figure S1.**
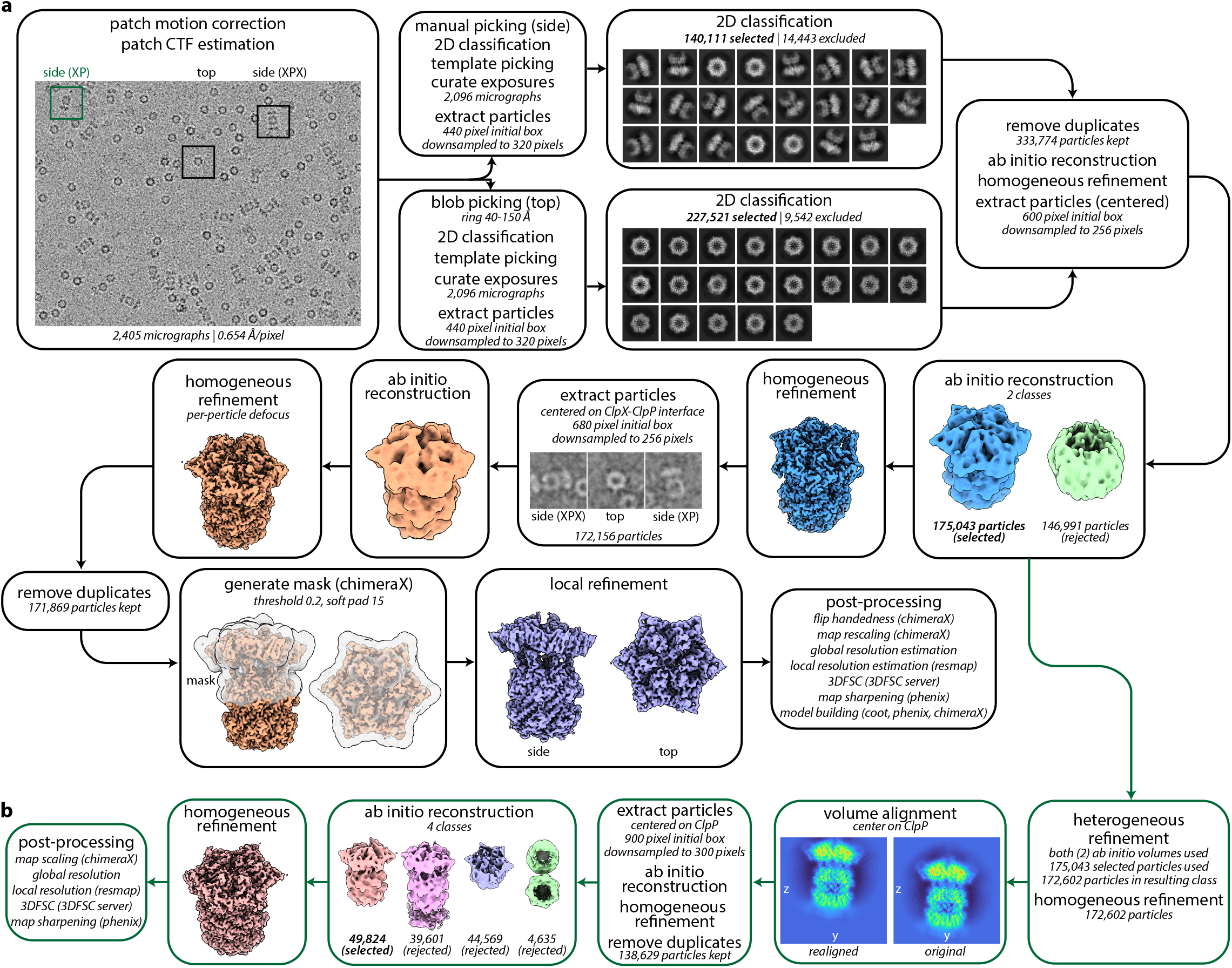
CryoSPARC processing workflow for ^SC^ClpX^ΔN^•ClpP particles. (**a**) ^SC^ClpX^ΔN^•ClpP structures derived from mixtures of singly and doubly capped particles. (**b**) ^SC^ClpX^ΔN^•ClpP from specifically singly capped particles. Note that the singly capped particles are an extracted subset of the full particle stack. Job names, job details, and non-default parameters (italicized) noted in each box.

**Figure S2.**
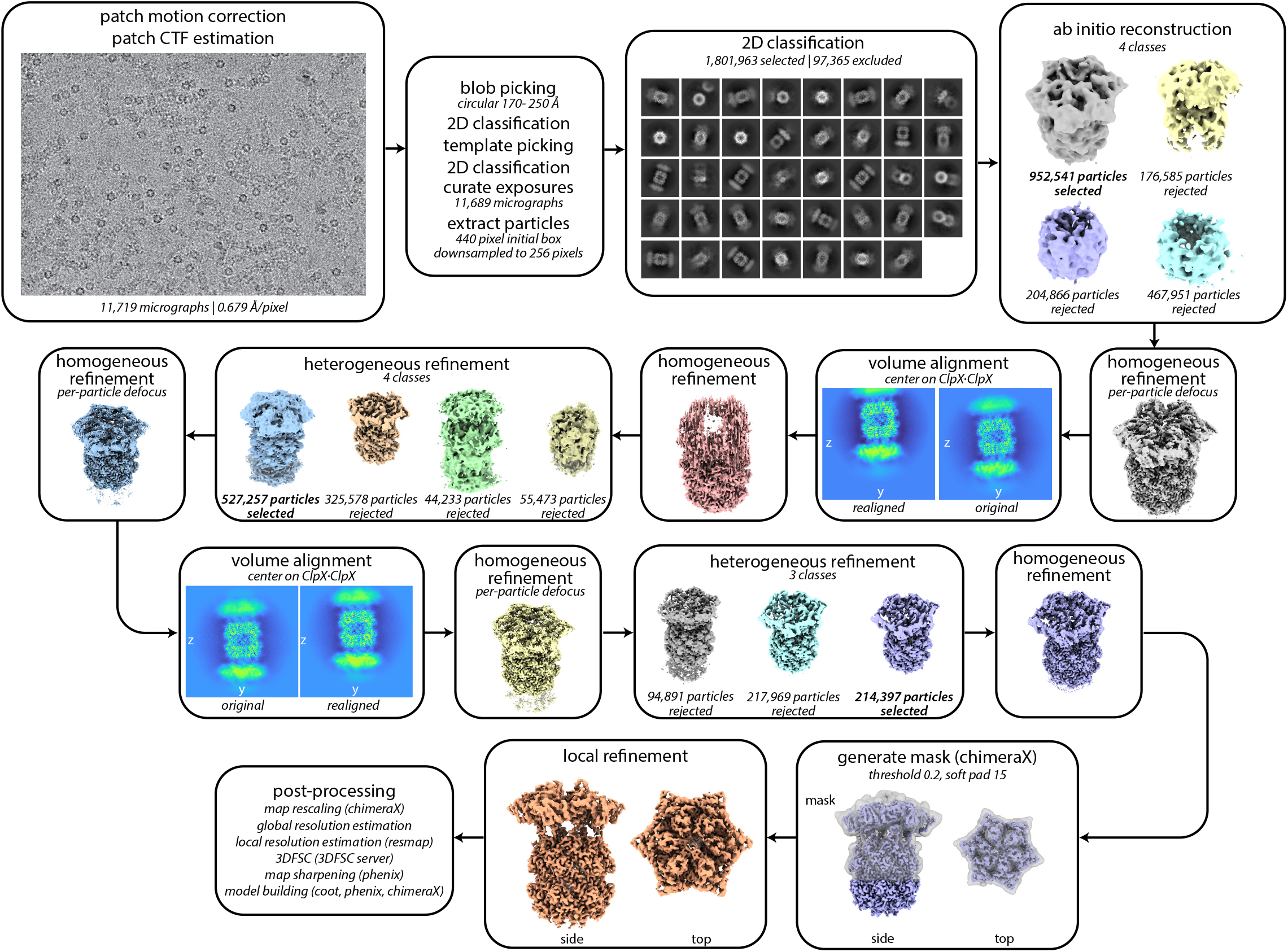
CryoSPARC processing workflow for full-length ClpX•ClpP particles. Job names, job details, and non-default parameters (italicized) are noted in each box.

**Figure S3.**
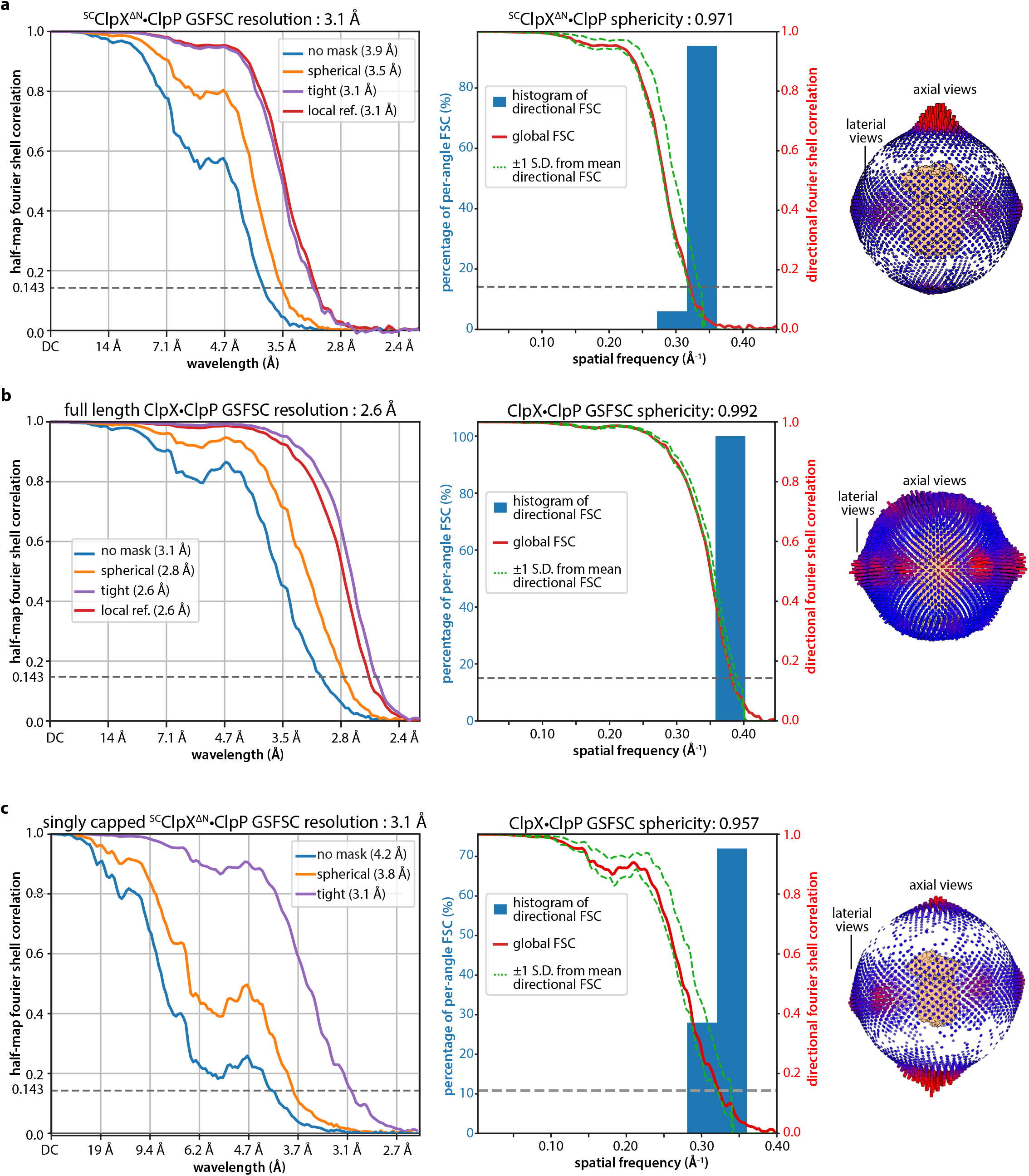
Global resolution, directional resolution, and projection angle distribution. (**a**) ^SC^ClpX^ΔN^•ClpP. (**b**) Full-length ClpXP. (**c**) ^SC^ClpX^ΔN^•ClpP from singly capped particles.

**Figure S4.**
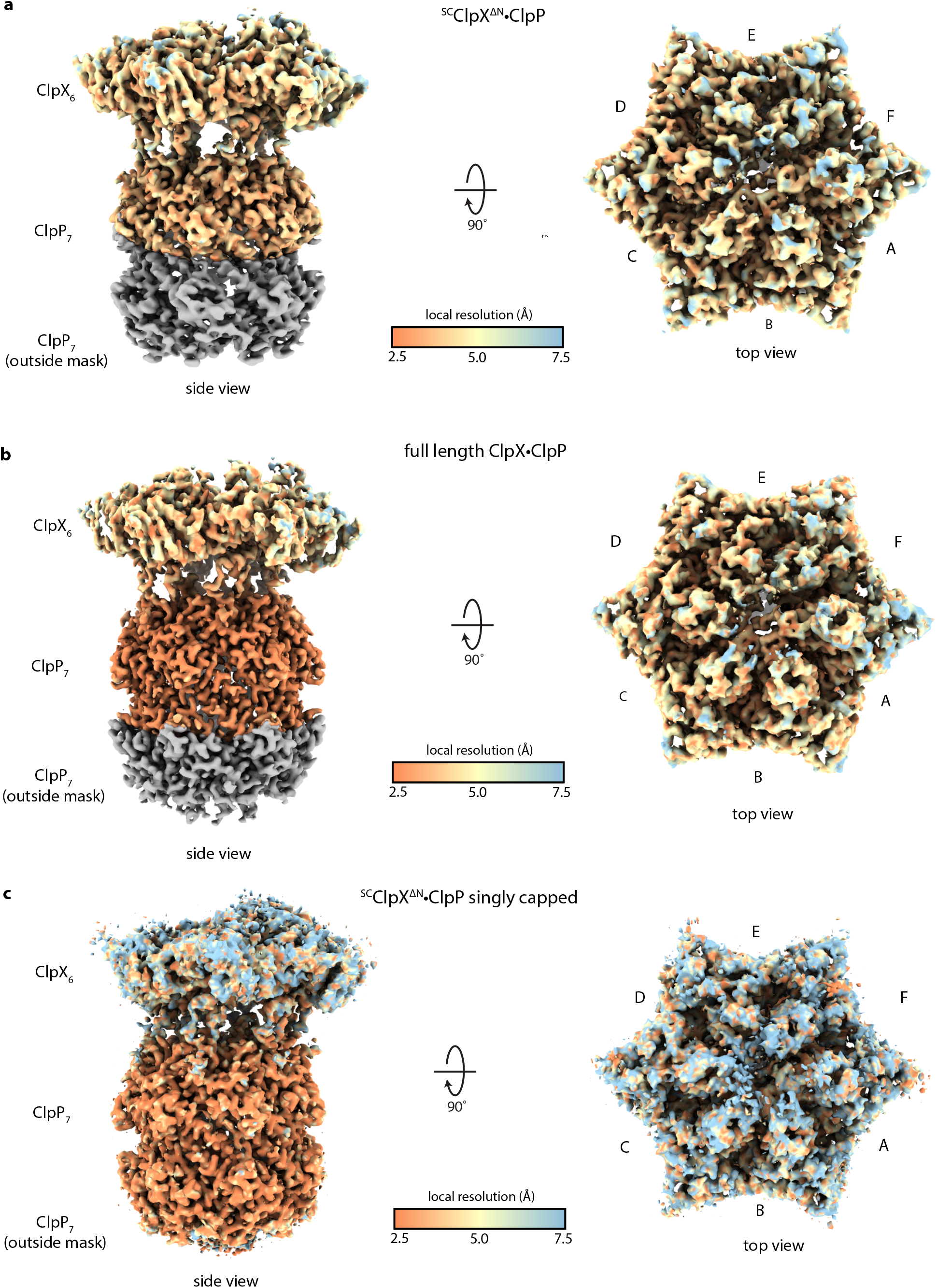
Local resolution assessment. Maps colored by local resolution as estimated by the cryoSPARC implementation of monoRes^46^. Regions outside of the mask used for local refinement in panels (a) and (b) are colored gray. (**a**) Side and top views of ^SC^ClpX^ΔN^•ClpP. (**b**) Side and top views of full-length ClpXP. (**c**) Side and top views of ^SC^ClpX^ΔN^•ClpP from singly capped particles.

**Figure S5.**
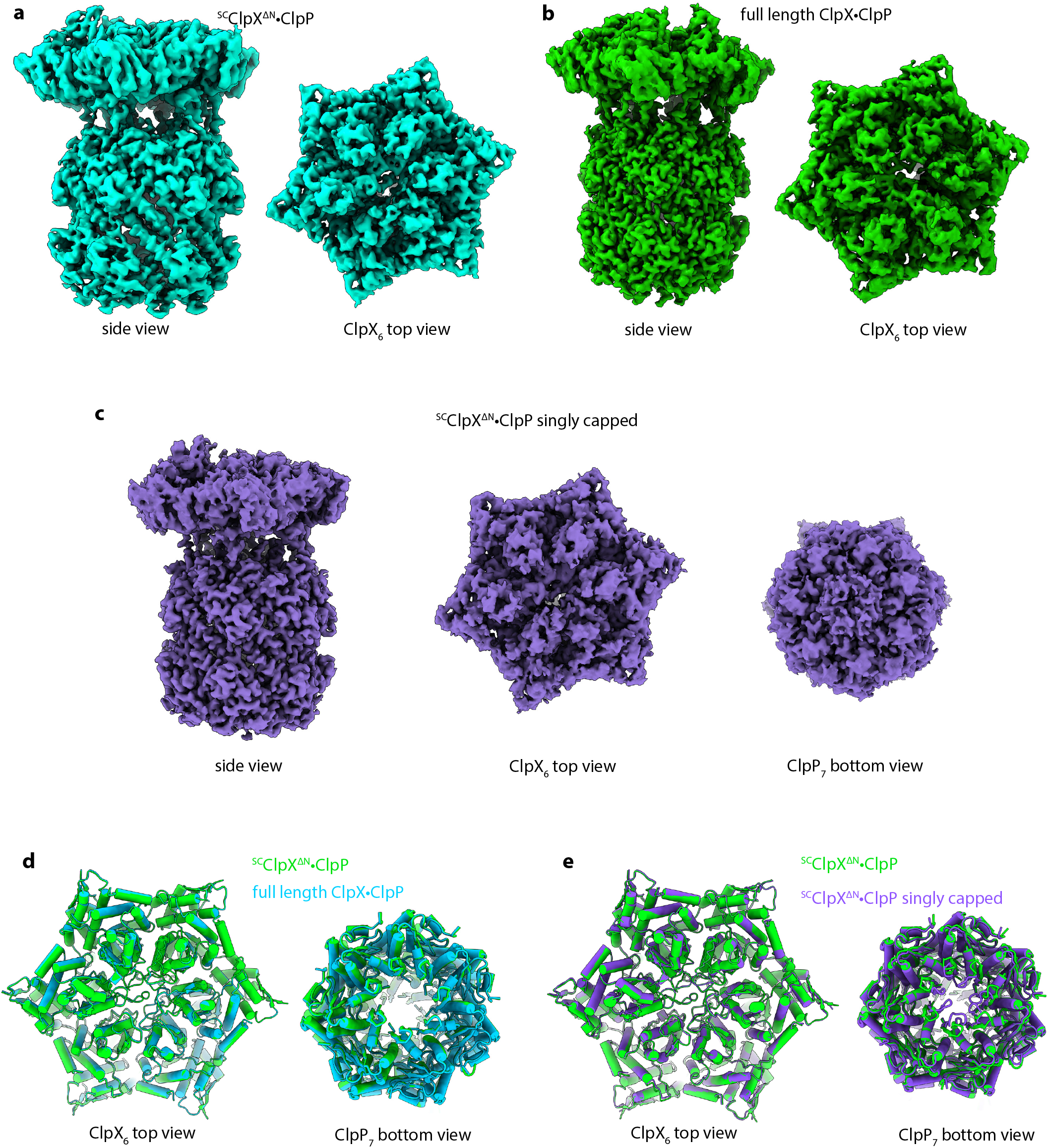
Density maps and models. (**a**) Cryo-EM density of ^SC^ClpX^ΔN^•ClpP in side and top views. (**b**) Cryo-EM density of full-length ClpXP in side and topviews. (**c**). Cryo-EM density of singly capped ^SC^ClpX^ΔN^•ClpP in side, top, and bottom views. (**d**). Alignment of full-length ClpXP (cyan; cartoon representation) and ^SC^ClpX^ΔN^•ClpP atomic models (green; cartoon representation) has a Cα RMSD of ∼0.3 Å. (**e**) Alignment of ^SC^ClpX^ΔN^•ClpP (green; cartoon representation) and exclusively singly capped particles (purple; cartoon representation) has a Cα RMSD of ∼0.9 Å. Note the axial portal of the distal Clp ring is closed in the exclusive singly capped ^SC^ClpX^ΔN^•ClpP particles (c, purple model in e) and open in the predominantly doubly capped ^SC^ClpX^ΔN^•ClpP particles (green model in e).

**Table S1.** Cryo-EM data collection, processing, model building, and validation statistics.

| Sample and data deposition information |  |  |  |
| --- | --- | --- | --- |
| Sample name | Full-length ClpX•ClpP | <sup>SC</sup>ClpX<sup>ΔN</sup>•ClpP | Singly capped <sup>SC</sup>ClpX<sup>ΔN</sup>•ClpP |
| Nucleotide added | ATP | ATPyS | ATPyS |
| PDB ID | 8E91 | 8E8Q | 8E7V |
| EMDB ID | 27952 | 27946 | 27941 |
| EMPIAR ID | to be deposited | to be deposited |  |
| Data collection |  |  |  |
| Microscope | Titan Krios G3i |  |  |
| Camera | Gatan K3 direct detection camera (counting mode) |  |  |
| Magnification (nominal) | 130,000 X | 105,000 X |  |
| Accelerating voltage (kV) | 300 |  |  |
| Total electron dose (e<sup>-</sup>/Å<sup>2</sup>) | 52.7 | 49.4 |  |
| Defocus range (μm) | -0.3 to -1.75 | -0.75 to -2.5 |  |
| Micrographs collected | 11,719 | 2,405 |  |
| Pixel size (Å) initial/calibrated | 0.679 / 0.654 | 0.435 / 0.416 |  |
| Map reconstruction |  |  |  |
| Image processing package | cryoSPARC<sup>44</sup> |  |  |
| Total extracted particles | 1,899,328 | 367,632 |  |
| Final particle count | 214,397 | 171,869 | 49,824 |
| Symmetry imposed | C1 |  |  |
| Resolution (Å) |  |  |  |
| 0.143 GSFSC unmasked | 3.1 | 3.9 | 4.2 |
| 0.143 GSFSC spherical mask | 2.8 | 3.5 | 3.8 |
| 0.143 GSFSC tight/local mask | 2.6 | 3.1 | 3.1 |
| 3DFSC sphericity (out of 1.0) | 0.992 | 0.971 | 0.957 |
| Model composition |  |  |  |
| Non-hydrogen atoms | 70,125 | 70,103 | 70,085 |
| Protein residues | 4,783 | 4,782 | 4,779 |
| Ligands | 9 | 9 | 9 |
| Model refinement |  |  |  |
| Refinement package | Phenix<sup>45</sup> |  |  |
| Map-to-model cross correlation | 0.83 | 0.80 | 0.82 |
| RMS deviations bond lengths (Å) | 0.003 | 0.003 | 0.004 |
| RMS deviations bond angles (°) | 0.650 | 0.625 | 0.665 |
| Model validation |  |  |  |
| MolProbity score | 0.91 | 0.87 | 0.83 |
| Clash score | 1.63 | 1.40 | 1.18 |
| C-beta outliers (%) | 0 | 0 | 0 |
| Rotamer outliers (%) | 0 | 0 | 0 |
| Ramachandran favored (%) | 99.2 | 99.2 | 99.4 |

## Notes

### Competing Interest Statement

The authors have declared no competing interest.

